# A novel binary *k*-mer approach for classification of coding and non-coding RNAs across diverse species

**DOI:** 10.1101/2021.06.21.449245

**Authors:** Neha Periwal, Priya Sharma, Pooja Arora, Saurabh Pandey, Baljeet Kaur, Vikas Sood

## Abstract

Classification among coding (CDS) and non-coding RNA (ncRNA) sequences is a challenge and several machine learning models have been developed for the same. Since the frequency of curated coding sequences is many-folds as compared to that of the ncRNAs, we devised a novel approach to work with the complete datasets from fifteen diverse species. In our proposed novel binary approach, we replaced all the ‘A’s and ‘T’s with ‘0’s and ‘G’s and ‘C’s with ‘1’s to obtain a binary form of coding and ncRNAs. The *k*-mer analysis of these binary sequences revealed that the frequency of binary patterns among the coding and ncRNAs can be used as features to distinguish among them. Using insights from these distinguishing frequencies, we used *k*-nearest neighbour classifier to classify among them. Our strategy is not only time-efficient but leads to significantly increased performance metrics including Matthews correlation coefficient (MCC) for some species like *P. paniscus, M. mulatta, M. lucifugus, G. gallus, C. japonica, C. abingdonii, A. carolinensis, D. melanogaster* and *C. elegans* when compared with the conventional ATGC approach. Additionally, we also show that the values of MCC obtained for diverse species tested on the model based on *H. sapiens* correlated with the geological evolutionary timeline thereby further strengthening our approach. Therefore, we propose that CDS and ncRNAs can be efficiently classified using “2-character” frequency as compared to “4-character” frequency of ATGC approach. Thus, our highly efficient binary approach can replace the more complex ATGC approach successfully.

## INTRODUCTION

The biological information is decoded in the macromolecule known as the Deoxyribonucleic Acid (DNA). The information from the DNA is translated into proteins via intermediary macromolecules commonly known as the Ribonucleic Acid (RNA). RNA molecules can be categorized as either coding or non-coding. Recently, non-coding RNA (ncRNA) has shown to play a vital role in various biological processes [1]. The complexity of ncRNAs can be conjectured by the fact that the human genome encodes for more than one million ncRNAs which is many folds as compared to the number of coding sequences (CDS) [2]. Apart from its abundance, the repertoire of ncRNA has been directly related to the biological complexity [3]. Non-coding transcripts have been shown to play a critical role in almost all the biological processes. The sheer diversity of the role that they play among biological processes ranges from regulating various biological phenomena including reproduction [4], embryonic development [5], immune responses [6], cancer [7] to viral infections [8–10]. Apart from their indispensable role in biological processes, several ncRNAs have also been shown to act as biomarkers in several diseases including cancer, tuberculosis, dengue and autoimmune diseases [11–17].

Owing to their role and propensity towards disease diagnosis and pathogenesis, categorization and characterization of ncRNAs is essential to understand the fundamentals of biological processes and disease progression. Although there are well characterized features including open reading frame (ORF) and protein conservation for coding transcripts which could be used for differentiating coding from non-coding transcripts, identification and classification of non-coding transcripts is still a big challenge [18,19]. Additionally, next generation sequencing is generating huge amounts of data consisting of thousands of novel transcripts thereby making ncRNA identification and classification a mammoth task. Therefore, immediate steps are required to devise novel methods that can classify CDS and ncRNAs with high accuracy and reliability. In this direction, several groups have successfully used machine learning (ML) based approaches which have significantly contributed towards the ncRNA identification and classification. These models are primarily dependent on features which are derived from transcript nucleotide sequences using mathematical approaches [20]. For example, the support vector machine (SVM) classifier based Coding Potential Calculator (CPC) utilised six features of transcripts to classify 5,610 CDS and 2,670 ncRNAs with good accuracy [21]. Another SVM model by Sun et al. utilized ten features from the broad categories including sequence conservation, ORF features and frequencies of seven di-nucleotide or tri-nucleotide sequences to identify long intergenic non-coding RNA (lincRNA) sequences. The study included 5,177 and 889 randomly selected sequences from human and mouse respectively and demonstrated higher prediction accuracy [22]. However, one of the drawbacks of the model was that it was restricted to sequences obtained from human and mouse only. Another tool named Coding Non-Coding Index (CNCI) was created that was based on intrinsic sequence information i.e., frequency of adjoining nucleotide triplets (ANT) to predict Most-Like CDS regions (MLCDS). Although the model was able to predict ncRNAs in a variety of species especially with poor genomic annotation, it had the tendency to misclassify transcripts with sequencing errors such as insertions and deletions [23]. The shortcomings of the CNCI tool were taken care of by another binary classifier named PLEK (predictors of long-non coding RNAs and messenger RNA) which was based on an improved *k*-mer scheme and SVM algorithm [24]. The classifier was trained on 34,691 human CDS and 22,389 ncRNA sequences and provided better specificity and sensitivity as compared to CNCI on real sequencing data. Another tool named long non-coding RNA identification tool (LncRNA-ID) was built on random forest (RF) classification method [25] and explored the coding potential of a transcript based on multiple features broadly covering three categories i.e., ORF, ribosome interaction and protein conservation. Other SVM based method utilized various features derived from gene structure, transcript sequence, potential codon sequence, conservation, ORF length, ORF relative length and frequencies of nucleotide pattern to improve the sensitivity and accuracy of CDS and ncRNAs from humans and several other species [26]. Studies mentioned above used ML models involving several features based on ORFs or protein conservation to characterize between CDS and ncRNAs. However, features based on ORFs or protein conservation might be inept owing to the fact that certain ncRNAs especially lncRNAs may have long putative ORFs or evolutionary conserved sequences [27]. Moreover, certain coding transcripts don’t have conserved ORFs [28] resulting in high false positive and false negative findings.

In order to attain more specificity and sensitivity, several groups have successfully utilized *k*-mer based deep learning approaches for classification and characterization of CDS and ncRNAs. In one of the studies, it was observed that ncRNAs performing related functions had a similar *k*-mer profile. This observation enabled them to cluster ncRNAs based on their functional profile [29]. Another study used *k*-mer embedding vectors to construct a deep-learning model to distinguish among CDS and ncRNA sequences. This five layered model included Bidirectional Long Short Term Memory (BLSTM) layer, a convolutional neuronal network (CNN) layer and three hidden layers to achieve 96.4% accuracy (30). In one study, *k*-mer approach was used to differentiate among CDS and ncRNAs [31] and was also used for classification of ncRNAs into relevant families in a different study [32].

Majorly, the experiments and results reported used balanced training and testing datasets that are devised to handle the uneven number of the positive and negative class samples. Although *k*-mer based approaches have been explored in several models including deep-learning, these approaches are generally computationally intensive. The complexity of the searches based on the four nucleotides is high. Since DNA consists of four nucleotides ‘A’, ‘T’, ‘G’ and ‘C’, there are 16 unique combinations for a *2*-mer, 64 combinations for *3*-mer and 256 combinations for *4-mer* that need to be investigated. As the size of *k* grows, it can be observed that the number of combinations grows exponentially, hence the volume of search becomes complex.

In order to have a robust feature set and to overcome the pertinent challenge of time complexity, in this paper, we have devised the binary *k*-mer approach to differentiate among CDS and ncRNAs. In this approach the ‘A’s and ‘T’s nucleotides are replaced with a zero ‘0’ whereas ‘G’s and ‘C’s with a one ‘1’. In essence the nucleotide sequences that were made of the symbols ‘A’, ‘T’, ‘G’ and ‘C’ are replaced with sequences of 0s and 1s. This approach was formulated keeping in mind that ‘A’ and ‘T’ both pair with each other with double bonds (low energy) whereas ‘G’ and ‘C’ pair with each other using triple bonds (high energy). The ML models were then built to classify CDS and ncRNAs using the features extracted from the binary approach. In order to validate our binary approach, exhaustive experiments were carried out on fifteen phylogenetically diverse species including *Homo sapiens, Pan paniscus, Macaca mulatta, Mus musculus, Myotis lucifugus, Gallus gallus, Coturnix japonica, Chelonoidis abingdonii, Anolis carolinensis, Xenopus tropicalis, Leptobrachium leishanense, Salmo trutta, Danio rerio, Drosophila melanogaster and Caenorhabditis elegans.* The *k*-mer analysis of these binary genomes revealed the enrichment of distinguishing patterns among CDS and ncRNAs among all the species. Using the frequency of various *k*-mer of each sequence in binary files as the features, we show that our ML model based on *k*-nearest neighbour (*k*NN) is highly efficient in classification of CDS and ncRNAs as compared to the traditional ATGC *k*-mer approach both in terms of computational time as well as Matthews correlation coefficient (MCC) which is the preferred metric to evaluate the performance of the imbalanced data. Thus, we have identified a unique binary strategy which can be explored to differentiate among CDS and ncRNAs in a highly efficient manner.

## MATERIAL AND METHODS

### Coding and ncRNA datasets

The CDS as well as ncRNA sequences for all the fifteen species included in this study were manually downloaded from Ensembl (https://asia.ensembl.org/info/data/ftp/index.html) in FASTA format on 30 October 2020 [33]. In order to capture the genetic diversity, we included the species on the following criteria: (a) both coding and ncRNA data should be available for all the species and (b) the species should have been extensively characterized. Based on the above-mentioned criteria’s, representative sequences from various diverse species from important classes including mammals, aves, reptiles, amphibians, pisces, insects and nematodes were included. In this study, we mainly focused on the model and highly characterized organisms including *Homo sapiens, Pan paniscus, Macaca mulatta, Mus musculus, Myotis lucifugus, Gallus gallus, Coturnix japonica, Chelonoidis abingdonii, Anolis carolinensis, Xenopus tropicalis, Leptobrachium leishanense, Salmo trutta, Danio rerio, Drosophila melanogaster and Caenorhabditis elegans*.

### Pre-processing of data

#### Removal of characters other than ‘A’, ‘T’, ‘G’ and ‘C’

In order to make sure that the input files did not contain any character other than ‘A’, ‘T’, ‘G’, and ‘C’, the input file (in FASTA format) was processed using python script that removes all the characters other than ‘A’, ‘T’, ‘G’, and ‘C’. The script was run on both the coding as well as ncRNA input files of all the species.

#### Removal of short sequences

Once the characters other than ‘A’, ‘T’, ‘G’, and ‘C’ were removed from the input file, we removed short sequences from both the files. We retained all the sequences greater than 50 nucleotides in our coding and ncRNA files so as to include them in our study.

### Conversion of DNA sequences into binary form

Several groups have investigated CDS and ncRNA sequences using conventional approaches taking all the four nucleotides into account which thereby leads to enhanced complexity. However, we speculated that we could replicate their key findings using only two characters (’0’s and ‘1’s) representing all the four nucleotides. Therefore, all the ‘A’s and ‘T’s were replaced by ‘0’s whereas ‘G’s and ‘C’s were replaced by ‘1’s in both the coding and ncRNA processed files of all the fifteen species. This leads to the generation of binary files where each sequence is represented by a string of ‘0’s and ‘1’s. This replacement decreased the genome complexity as well as requirement of computer hardware to process the full genomic sequences.

### Generation of possible ***k-***mer patterns

The next step towards the analysis of coding and ncRNA binary files involved generating all the possible patterns for a given *k*-mer. For each *k*-mer, 2^k^ combinations of patterns were extracted for each file.

### Finding frequency of binary ***k***-mer in CDS and ncRNAs

For every 2^k^ pattern of a given *k*-mer (*k =* 3 to *k =* 18), sequences were read in an overlapping manner to calculate the corresponding frequency of that pattern in a binary CDS and ncRNA files. All the steps of data pre-processing, binary conversion and calculating frequencies were performed by inhouse python script.

### Machine learning model

We used the *k*-Nearest Neighbour (*k*NN), a well-known classifier to report the performance metrics of the experiments. Each sequence of the binary file of coding and ncRNA, was read in an overlapping manner to calculate the frequency for a given *k*-mer. The input file contains the number of sequences along the rows and the frequency of each pattern corresponding to the sequence along the columns. For all the species, the input file was prepared using the python inhouse script. Our next important task was to find the most suitable *k* (for the *k*-mer) to distinguish CDS from ncRNAs. To determine the optimum *k*-mer, we tested the *k*NN (with 5 neighbours) using the input file at different *k*-mer (*k =* 3 to *k* = 10). The value of *k* which gave the best performance metrics was chosen as the optimum *k* for further experiments.

#### Performance Measures

For each experiment, 10 independent runs of 10-fold cross validation were performed for the *k*NN classifier, with 5 neighbours. By averaging over the ten runs of the 10-fold cross validation, a realistic performance score was reported for all our experiments. For each experiment, True Positives (TP) is the number of correctly classified coding RNAs, True Negatives (TN) is the number of correctly classified ncRNAs, False Negatives (FN) is the number of CDS predicted as ncRNAs and False Positives (FP) is the ncRNAs predicted as CDSs. The performance of the ML algorithm was quantified in terms of classification accuracy, precision, recall, F1 score, area under the curve-receiver operating characteristics (AUC-ROC) and MCC as described.

i. Classification accuracy: Classification accuracy is the ratio of the correctly classified sequences.

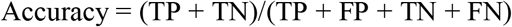
ii. Precision is the ratio of correctly classified CDS in the set of True and False Positives.

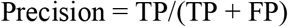
iii. Recall (sensitivity): Recall is the proportion of the correctly classified CDS in the set of all available coding sequences.

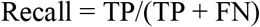
iv. F1 score: F1 score is the harmonic mean of precision and recall

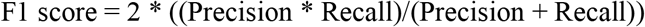
v. AUC-ROC: The AUC-ROC is commonly used to compare classification models for imbalanced problems.
vi. Matthews correlation coefficient (MCC): MCC is a reliable statistical metric which gives a high score when most of the positive and the negative predictions are correct.

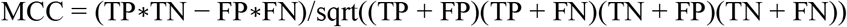

#### Optimum k-mer for high classification performance

Our experiments used all the available data of CDS and ncRNAs to build our model using 10 independent runs of 10-fold cross validation. The average results of each performance measure are reported. Through these experiments we found the optimum *k*-mer for classifying the CDS and ncRNAs across the species.

#### Efficacy of the binary approach in comparison to the ATGC k-mer approach

Further, to establish that we do not lose any relevant information by converting the ‘A’, ‘T’ ‘G’ and ‘C’ sequences into binary sequences, we compared the performance of the ATGC as well the binary *5*-mer approach for all the species included in this study. The reason why we have used binary *5*-mer is due to its best average performance across all species. The results of the MCC measures as well as the time taken for building the feature frequency datasets and the average model building time for all the species for *k =* 5 are reported. These experiments are conducted with the *k*NN classifier with ten independent runs of 10-fold cross validations.

#### Evaluation of the model on novel species

There is a need to classify the CDS and ncRNA for all possible species, but in the case of some species, due to lack of sufficient and relevant data, ML models cannot be built. The evolutionary relationship among species motivated us for our next set of experiments. The *k*NN model was built with the *5*-mer patterns as the features on the human dataset. The model was evaluated across several new species which were not part of this study. The MCC scores are also depicted on the time tree which was built using Time Tree web server (34).

## RESULTS AND DISCUSSION

### Identification of highly occurring patterns among human CDS

As the first line of study, we used CDS of the human genome to investigate whether any binary pattern exists across the sequences. We used computational approaches to identify patterns among the human CDS. We downloaded the human CDS sequences (in FASTA format) from Ensembl. We then performed all the pre-processing steps and converted the CDS file into the binary form by replacing ‘A’s, ‘T’s with ‘0’s and ‘G’s, ‘C’s with ‘1’s as described above. The resulting human CDS file now consisted of ‘0’s and ‘1’s only. We started our investigation with the set of *3*-mer patterns. In order to find the most commonly occurring patterns, we wrote a python script to calculate the frequency of all the possible *3*-mer (of ‘0’ and ‘1’) in the human coding sequences. Our analysis revealed that out of all the possible eight combinations, the pattern ‘101’ has the highest occurrence followed by the pattern ‘010’ and ‘011’. The pattern ‘000’ has the lowest occurrence (Figure 1A, Supplementary Table S1). This data suggested that binary human CDS had distinct frequency of patterns. It was also observed that the frequency of the highest occurring pattern ‘101’ was 1.74 times higher as compared to the lowest occurring one ‘000’. This difference in the frequency of various patterns among the binary coding sequences pointed towards a possible ‘binary signature’ among human CDS and encouraged us to take the approach further. We then investigated human CDS by calculating occurrences of all the possible combinations of *4*-mer, *5*-mer, and so on, upto *18*-mer. We observed that the ratio of the highest occurring pattern to the lowest occurring pattern kept on increasing with the increase in the *k*-mer length (Figure 1B). A closer inspection of top occurring sequences revealed the repetitive occurrence of the seed sequence i.e., ‘101’ that was obtained from the *3*-mer. We thus identified a stretch of ‘(101)_n_’ that was being enriched in human CDS (Figure 1C). This seed sequence is observed to be enriched up to *16*-mer in human CDS. However, in *17*-mer and *18*-mer we can observe that this seed sequence is no longer highly enriched but occupies a second position. Therefore our data suggest that there might be some binary patterns among the coding sequences.

**Figure 1.**
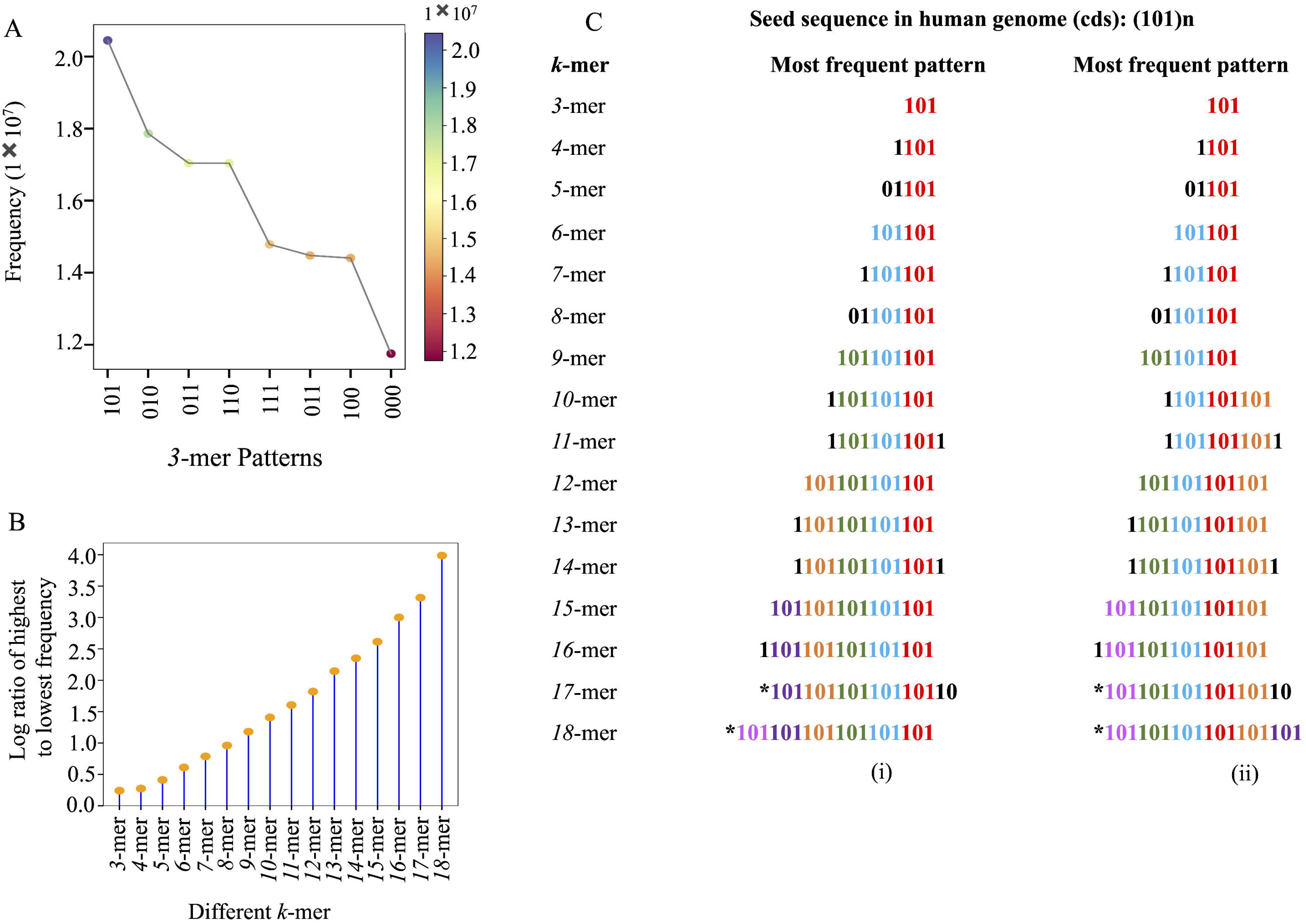
Identification of highly occurring patterns among human CDS (A) Pattern ‘101’ has the highest frequency while pattern ‘000’ has the lowest frequency among all *3*-mer combinations in human CDS. (B) The ratio of highest to lowest occurring pattern increases with the increase in the size of *k*-mer. (C) Possible arrangements of the highest occurring pattern among various *k*-mer (*k*=3 to 18) in human CDS. Note: ^*^ represents the second highest occurring pattern, (i) and (ii) represents possible different arrangements of highest occurring patterns.

### Frequent occurring pattern is the characteristic of CDS in several other species

Since analysis of the human CDS led us to the identification of the frequently occurring pattern, we sought to investigate whether we could replicate our findings among other species as well. Therefore, we used similar approach to investigate CDS of several other model and well characterized species including *Pan paniscus, Macaca mulatta, Mus musculus, Myotis lucifugus, Gallus gallus, Coturnix japonica, Chelonoidis abingdonii, Anolis carolinensis, Xenopus tropicalis, Leptobrachium leishanense, Salmo trutta, Danio rerio, Drosophila melanogaster and Caenorhabditis elegans.* The analysis of *3*-mer for several species, revealed that the pattern ‘101’ was the highest occurring in CDS for most of the species (Figure 2A). Similar results were obtained when the analysis was performed on *4*-mer and *5*-mer respectively (Figure 2B, 2C). As observed in humans, we found that the same patterns were enriched in most of the species. Moreover, the ratio of the highest occurring to the the lowest occurring pattern also increased with increase in the *k*-mer among all the species (Figure 2D). Therefore, we observed that a binary pattern (’101’) was highly enriched among

**Figure 2.**
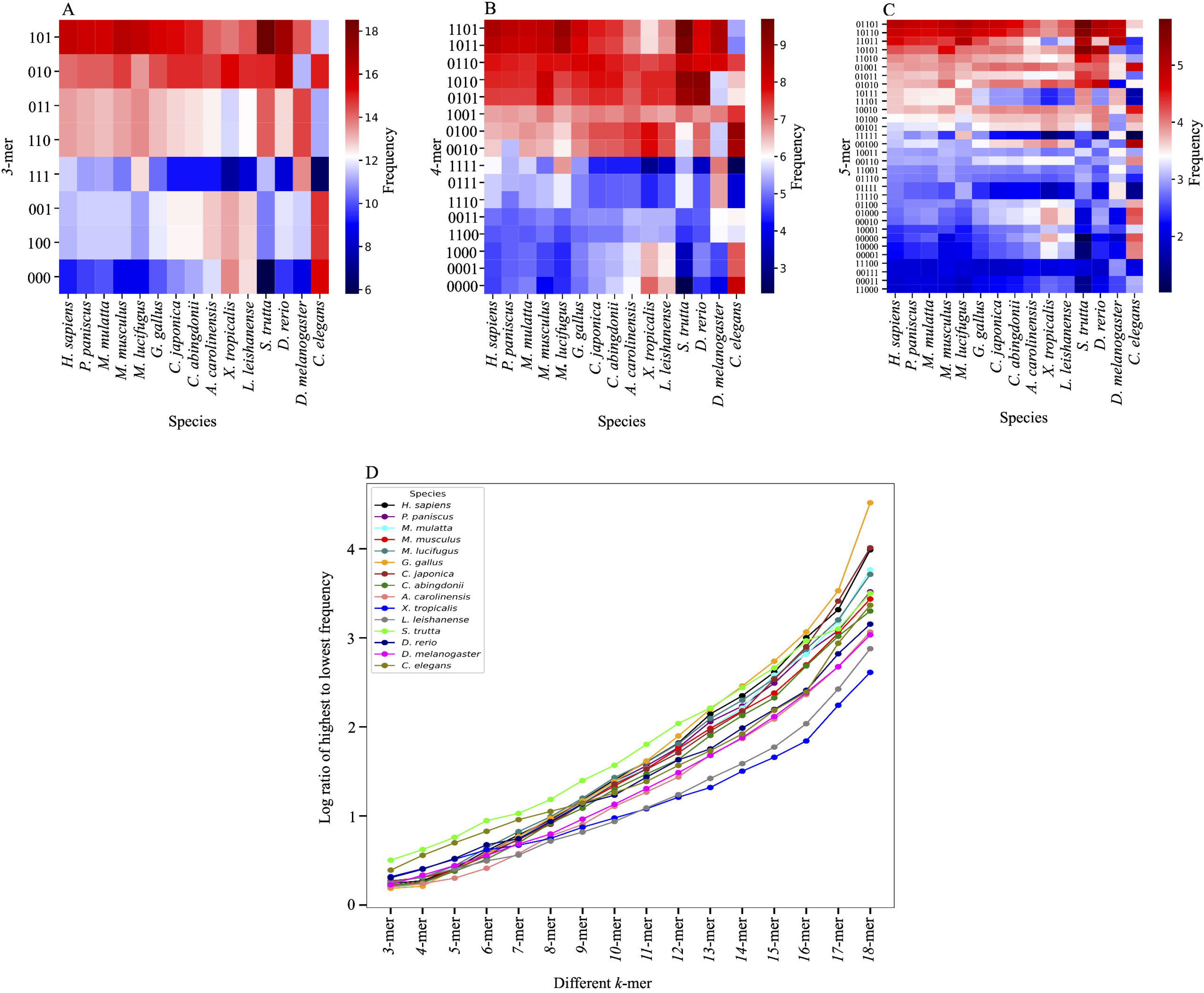
Frequent occurring pattern is the characteristic of CDS in several other species. Heatmap showing *k*-mer combinations and their frequencies in CDS of fifteen species (A) 3- mer combinations (B) 4-mer combinations (C) 5-mer combinations (D) Plot showing the ratio of highest to the lowest occurring pattern among CDS of fifteen species across different *k*-mer (k = 3 to k = 18). The ratio of highest to lowest occurring pattern increases with an increase in *k*-mer.

CDS of humans and other diverse species. Our results from CDS of various species suggested that there could be some conserved binary signatures among the CDS of species under the investigation. This data pointed towards some highly conserved binary signatures among the CDS of several species. The fact that similar sequences were enriched in multiple species pointed towards some role of these sequences in biological processes.

### Coding and non-coding sequences are characterized by unique occurring patterns

Since we have identified the highest occurring binary pattern among CDS of various species, we sought to investigate whether a similar pattern was enriched among the ncRNAs. Therefore, we planned to investigate ncRNAs of humans and all other species. Similar to the CDS, all the ncRNA sequences were obtained from Ensembl in a FASTA format and pre-processed in exactly the same way to obtain ncRNA binary files. We then performed the *k*-mer analysis on these ncRNA binary sequences to investigate whether there is an enrichment of similar pattern or not. We started our analysis with *3-*mer on human ncRNA binary sequences and observed that a particular pattern ‘000’ was highly enriched among them. (Figure 3A). We found that the highly enriched pattern ‘000’ in ncRNA was in fact the least occurring pattern in the human CDS (Figure 3B). This observation suggested that there was a huge difference among the architecture of human CDS and ncRNA sequences. Apart from the highly occurring sequences, it was also observed that frequency of various other binary patterns was different among CDS and ncRNA sequences. Moreover, similar to the CDS data, the ratio of the highest occurring to the lowest occurring pattern kept on increasing with an increase in the *k*-mer among human ncRNA binary sequences (Figure 3C). There was an enrichment of repeating pattern ‘(000)_n_’ among various *k*-mer in human ncRNA sequences (Figure 3D). The enrichment of seed sequence ‘000’ in case of ncRNA was different as compared to that of CDS where the enrichment of seed sequence ‘101’ was observed. We then investigated ncRNA binary data from the same the set of species that was used for CDS analysis and the same pattern ‘000’ was found to be enriched in most of them (Figure 3E). The frequency of binary patterns for various *k*-mer among CDS and ncRNAs was different and this observation helped us to formulate the strategy in distinguishing among coding and non-coding transcripts with high efficiency.

**Figure 3.**
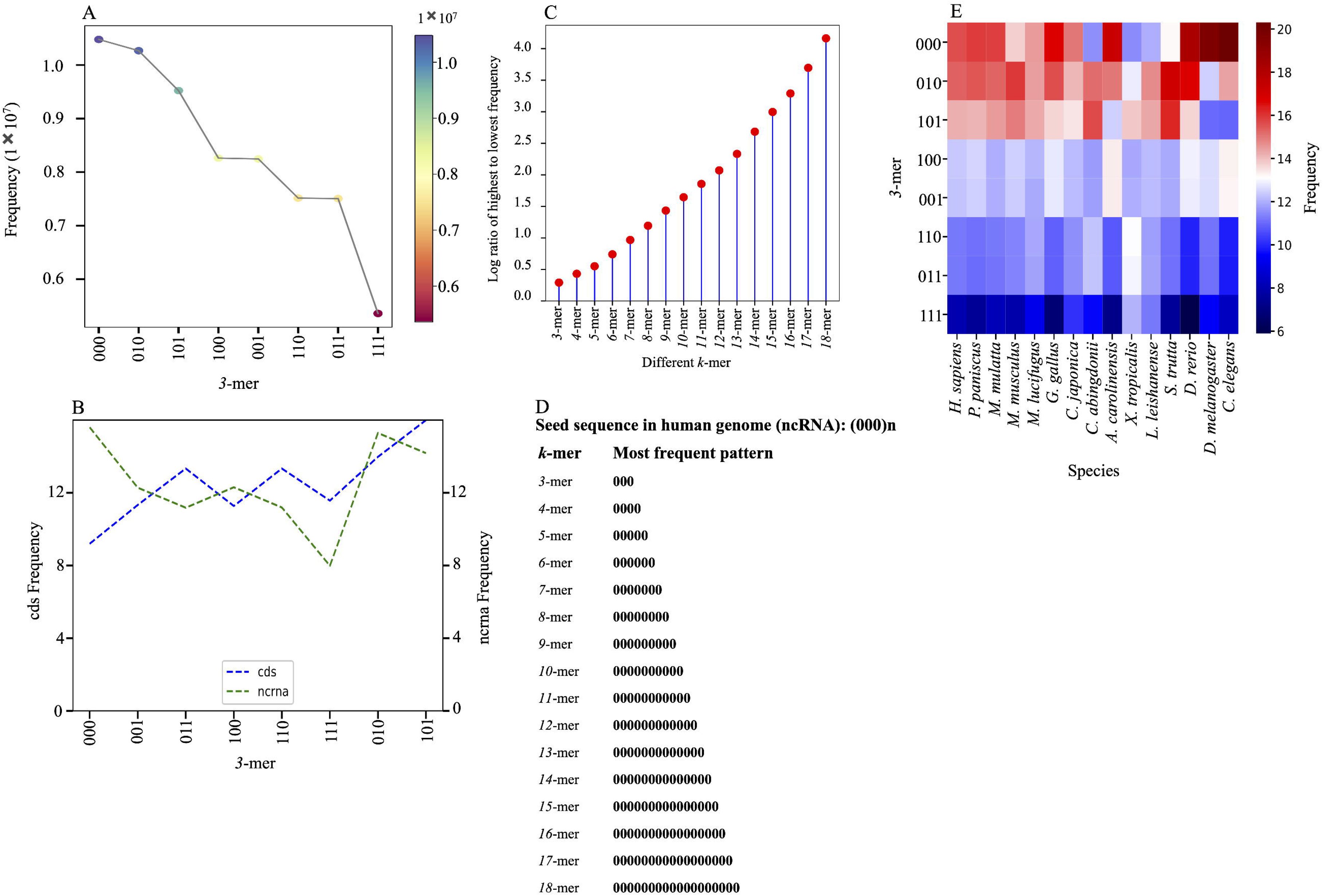
Coding and non-coding sequences are characterized by unique occurring patterns (A) Pattern ‘000’ has the highest frequency while pattern ‘111’ has the lowest frequency among all the 3-mer combinations in human ncRNA sequences. (B) Pattern ‘000’ has the highest frequency in human ncRNA whereas it is least occurring in human CDS. (C) The ratio of highest to lowest occurring pattern increases with an increase in the size of A>mer in human ncRNAs. (D) The seed sequence ‘(000)n’ increases with an increase in the *k*-mer for human ncRNAs and (E) Heatmap showing the pattern ‘000’ is enriched in most of the *3*-mer ncRNA sequences among fifteen species.

### Determining the optimum ***k***-mer for high classification performance

Extensive experiments have been performed to study the efficacy of the *k*NN classifier. The *k*NN classifier is suitable when there is no prior information available about the distribution of data and is a useful technique for non-linear data. Being a non-parametric classifier, it is used widely in ML problems. A thorough study of the optimum *k*-mer for various species was undertaken and the most suitable *k* value for the classifier was determined. Since the data of CDS and ncRNAs of all the species is highly skewed (Table 1), it becomes imperative to measure the performance in terms of metrics that take into consideration the imbalanced data problem. To adjudge the performance of classifiers in such biological data, measures like accuracy tend to give a high value even when the minority class gets heavily mis-classified. Other metrics that are reported are the TP, TN, precision and recall, which alone do not give the true efficacy of the classifier in an unbalanced setup. The metrics like AUC, F1 and MCC are used for classification and evaluation of imbalanced data. AUC is sensitive to class imbalance and does not have a closed form (35). The F1 score is commonly used for imbalanced data but it is independent of TN and is not symmetric for class swapping (36). In contrast, MCC generates a high score when the majority of positive data instances and the majority of negative data instances are correctly classified. MCC hence is effective as a relevant and correct measure of the efficiency of a decision algorithm. In our opinion, as illustrated mathematically and empirically in literature, MCC is the most appropriate measure to evaluate an unbalanced dataset (35, 36). We have used all the available data so as not to lose any significant information from the datasets. Some work in literature has randomly selected data to obtain balanced data using very less number of samples, and built machine learning models. This approach may be restrictive in nature as it does not include all possible data and hence the model is not trained as robustly. The high-performance measures reported might be unreliable in these studies. Owing to the preponderance of MCC towards the unbalanced data, we report the performance of the *k*NN classifier in terms of MCC in addition to other performance metrics (Supplementary Table S2).

**Table 1:** The number of CDS and ncRNAs from various species after the preprocessing step.

| S.No | Species | Number of Sequences |  |
| --- | --- | --- | --- |
|  |  | cds | ncRNA |
| 1 | *Homo sapiens* | 111199 | 58247 |
| 2 | *Pan paniscus* | 43225 | 9563 |
| 3 | *Macaca mulatta* | 48770 | 14665 |
| 4 | *Mus musculus* | 68026 | 22025 |
| 5 | *Myotis lucifugus* | 20714 | 4402 |
| 6 | *Gallus gallus* | 28442 | 10429 |
| 7 | *Coturnix japonica* | 27883 | 9577 |
| 8 | *Chelonoidis abingdonii* | 31797 | 2954 |
| 9 | *Anolis carolinensis* | 19175 | 7835 |
| 10 | *Xenopus tropicalis* | 54849 | 1384 |
| 11 | *Leptobranchium leishanense* | 49452 | 912 |
| 12 | *Salmo trutta* | 116557 | 5241 |
| 13 | *Danio rerio* | 52028 | 8113 |
| 14 | *Drosophila melanogaster* | 30580 | 3943 |
| 15 | *Caenorhabditis elegans* | 33484 | 9305 |

As depicted in Table 2, for *Homo sapiens*, the highest accuracy and MCC were obtained for *9*-mer. The performance measures for the fifteen species are recorded in Table 2 for *k =* 3 to *k =* 10. We can observe that the values of accuracy and MCC for all the other species were very encouraging. In case of *P. paniscus, M. mulatta, G. gallus, C. japonica, X. tropicalis,* and *D. melanogaster,* the highest MCC is observed for *5*-mer. For *M. lucifugus, C. abingdonii, A. carolinensis, and C. elegans, 4*-mer were the most distinguishing patterns. The species *M. musculus, L. leishanense, and D. reri* performed best for *7*-mer.

**Table 2:** The accuracy and MCC from *k =* 3 to *k =* 10 among various species. The bold numbers represent the highest value of MCC of a particular species. Average row represents the average of accuracy and MCC for the corresponding *k*-mer.

| Species | 3-mer |  | 4-mer |  | 5-mer |  | 6-mer |  | 7-mer |  | 8-mer |  | 9-mer |  | 10-mer |  |
| --- | --- | --- | --- | --- | --- | --- | --- | --- | --- | --- | --- | --- | --- | --- | --- | --- |
|  | Accuracy | MCC | Accuracy | MCC | Accuracy | MCC | Accuracy | MCC | Accuracy | MCC | Accuracy | MCC | Accuracy | MCC | Accuracy | MCC |
| *H. sapiens* | 0.795 | 0.535 | 0.826 | 0.607 | 0.858 | 0.681 | 0.6874 | 0.718 | 0.883 | 0.752 | 0.889 | 0.752 | 0.891 | **0.757** | 0.889 | 0.755 |
| *P. paniscus* | 0.881 | 0.554 | 0.895 | 0.607 | 0.897 | **0.620** | 0.889 | 0.582 | 0.880 | 0.542 | 0.872 | 0.497 | 0.864 | 0.455 | 0.857 | 0.418 |
| *M. mulatta* | 0.875 | 0.623 | 0.899 | 0.702 | 0.906 | **0.73** | 0.900 | 0.704 | 0.887 | 0.662 | 0.873 | 0.615 | 0.860 | 0.573 | 0.849 | 0.531 |
| *M. musculus* | 0.810 | 0.439 | 0.831 | 0.508 | 0.854 | 0.577 | 0.863 | 0.602 | 0.867 | **0.611** | 0.865 | 0.609 | 0.864 | 0.604 | 0.859 | 0.588 |
| *M. lucifugus* | 0.911 | 0.666 | 0.935 | **0.763** | 0.931 | 0.737 | 0.918 | 0.694 | 0.907 | 0.649 | 0.899 | 0.617 | 0.894 | 0.588 | 0.890 | 0.575 |
| *G. gallus* | 0.862 | 0.633 | 0.879 | 0.680 | 0.885 | **0.700** | 0.885 | 0.693 | 0.876 | 0.668 | 0.859 | 0.624 | 0.834 | 0.550 | 0.808 | 0.470 |
| *C. japonica* | 0.863 | 0.623 | 0.880 | 0.668 | 0.881 | **0.669** | 0.872 | 0.644 | 0.862 | 0.615 | 0.845 | 0.561 | 0.828 | 0.508 | 0.815 | 0.459 |
| *C. abingdonii* | 0.929 | 0.415 | 0.936 | **0.475** | 0.932 | 0.424 | 0.924 | 0.314 | 0.921 | 0.242 | 0.920 | 0.221 | 0.918 | 0.220 | 0.919 | 0.199 |
| *A. carolinensis* | 0.809 | 0.51 | 0.835 | **0.578** | 0.835 | 0.571 | 0.816 | 0.523 | 0.799 | 0.474 | 0.786 | 0.433 | 0.778 | 0.407 | 0.770 | 0.384 |
| *X. tropicalis* | 0.987 | 0.699 | 0.989 | 0.753 | 0.990 | **0.775** | 0.990 | 0.765 | 0.989 | 0.762 | 0.989 | 0.762 | 0.989 | 0.763 | 0.989 | 0.761 |
| *L. leishanense* | 0.987 | 0.560 | 0.989 | 0.668 | 0.992 | 0.728 | 0.992 | 0.742 | 0.992 | **0.752** | 0.992 | 0.743 | 0.992 | 0.747 | 0.992 | 0.745 |
| *S. trutta* | 0.973 | 0.630 | 0.976 | 0.675 | 0.978 | 0.711 | 0.979 | 0.718 | 0.979 | **0.717** | 0.979 | 0.717 | 0.979 | 0.717 | 0.979 | 0.713 |
| *D. rerio* | 0.905 | 0.542 | 0.925 | 0.647 | 0.934 | 0.692 | 0.937 | 0.704 | 0.938 | **0.711** | 0.935 | 0.697 | 0.934 | 0.687 | 0.931 | 0.670 |
| *D. melanogaster* | 0.943 | 0.699 | 0.949 | 0.725 | 0.949 | **0.728** | 0.949 | 0.728 | 0.948 | 0.711 | 0.944 | 0.692 | 0.941 | 0.676 | 0.937 | 0.646 |
| *C. elegans* | 0.905 | 0.706 | 0.914 | **0.732** | 0.903 | 0.700 | 0.879 | 0.618 | 0.867 | 0.535 | 0.841 | 0.469 | 0.834 | 0.429 | 0.829 | 0.407 |
| Average | **0.895** | **0.589** | **0.910** | **0.652** | **0.915** | **0.669** | **0.898** | **0.649** | **0.906** | **0.626** | **0.899** | **0.600** | **0.893** | **0.579** | **0.887** | **0.554** |

Using binary *k*-mer, we have substantially reduced the complexity of the determination of frequency as well as the building of the decision algorithm. To decide on the optimal value of *k*, the average rank over the 15 species was considered. On the basis of the average MCC among all the species that we studied, the efficacy of each *k*-mer was ranked in Figure 4A. It can be observed that on an average, the features formed using *5*-mer are the most distinguishing feature set for classifying CDS and ncRNAs. We can thus infer that ML models with binary *5*-mer are the most efficient for distinguishing CDS from the ncRNAs.

**Figure 4.**
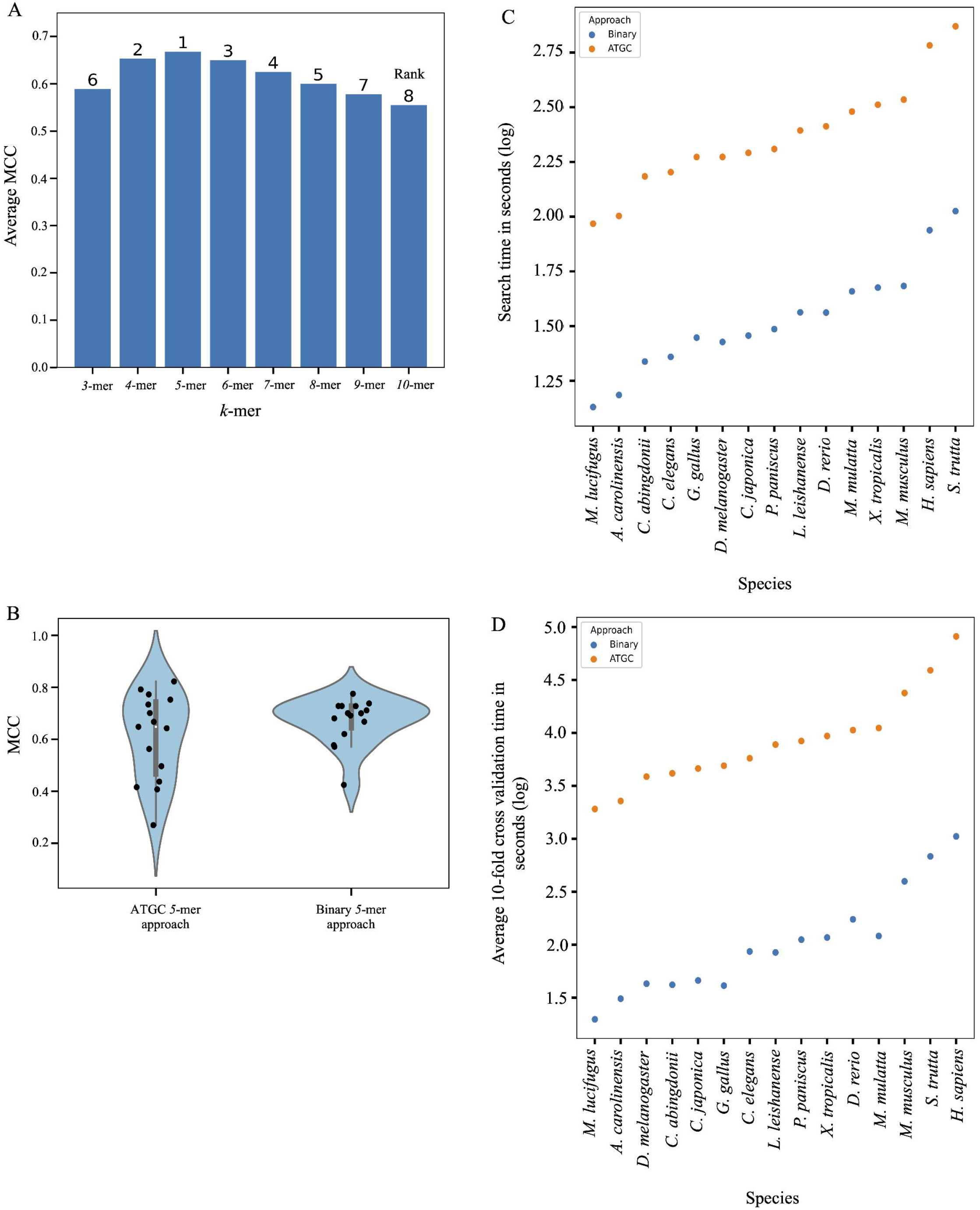
Comparison between ATGC and binary 5-mer approach (A) The ranking of average MCC score for different *k*-mer along all fifteen species using *k*NN classifier (B) showing the MCC scores (*k*NN with 5 neighbours) of fifteen species using ATGC and binary 5-mer approach (C) comparison of 5-mer search time taken by ATGC and binary approach for each species (D) comparison of 5-mer model building time taken by ATGC and binary approaches for each species.

### Comparison between ATGC and binary ***5***-mer approach

In literature, CDS and ncRNAs have been identified using the *k*-mer approach based on the ‘A’, ‘T’, ‘G’ and ‘C’ nucleotides. To gather the confidence in our binary *k*-mer approach, we illustrate the difference in the traditional ATGC and the proposed binary approach on the basis of performance metrics. We conducted experiments with the 1024 possible patterns for the traditional *5*-mer (ATGC approach) to classify the coding and non-coding sequences of the fifteen species, using the *k*NN classifier. This experiment was compared to the results of the binary *5*-mer approach with the 32 patterns only. This was to understand which one of the two approach is superior. Table 3 illustrates the MCC reported in both the cases. The average MCC of all the species in traditional ATGC *5*-mer was 0.608 whereas it was 0.669 in case of binary *5*-mer (Figure 4B). However, it can be seen that MCC values increased by nearly one and a half times of conventional ATGC approach in *G. gallus* and *C. elegans* thereby supporting the robustness of por binary approach. The time complexity is also tabulated in Table 3. The traditional approach took an average of 273.56 seconds to search for the *5*-mer patterns over fifteen species whereas it took 39.62 seconds to process the data among the same set of species using our proposed binary approach. Similarly, the traditional approach took an average of 14588.64 seconds to build the *k*NN model for the *5*-mer whereas the same model was built in 203.17 seconds using our binary approach (Figure 4C, 4D, Supplementary Table S3). This clearly suggests that binary *5*-mer with only 32 combinations out-performed the ATGC approach. In fact, the time complexity of the proposed approach is extremely less and the performance is highly encouraging.

**Table 3:**
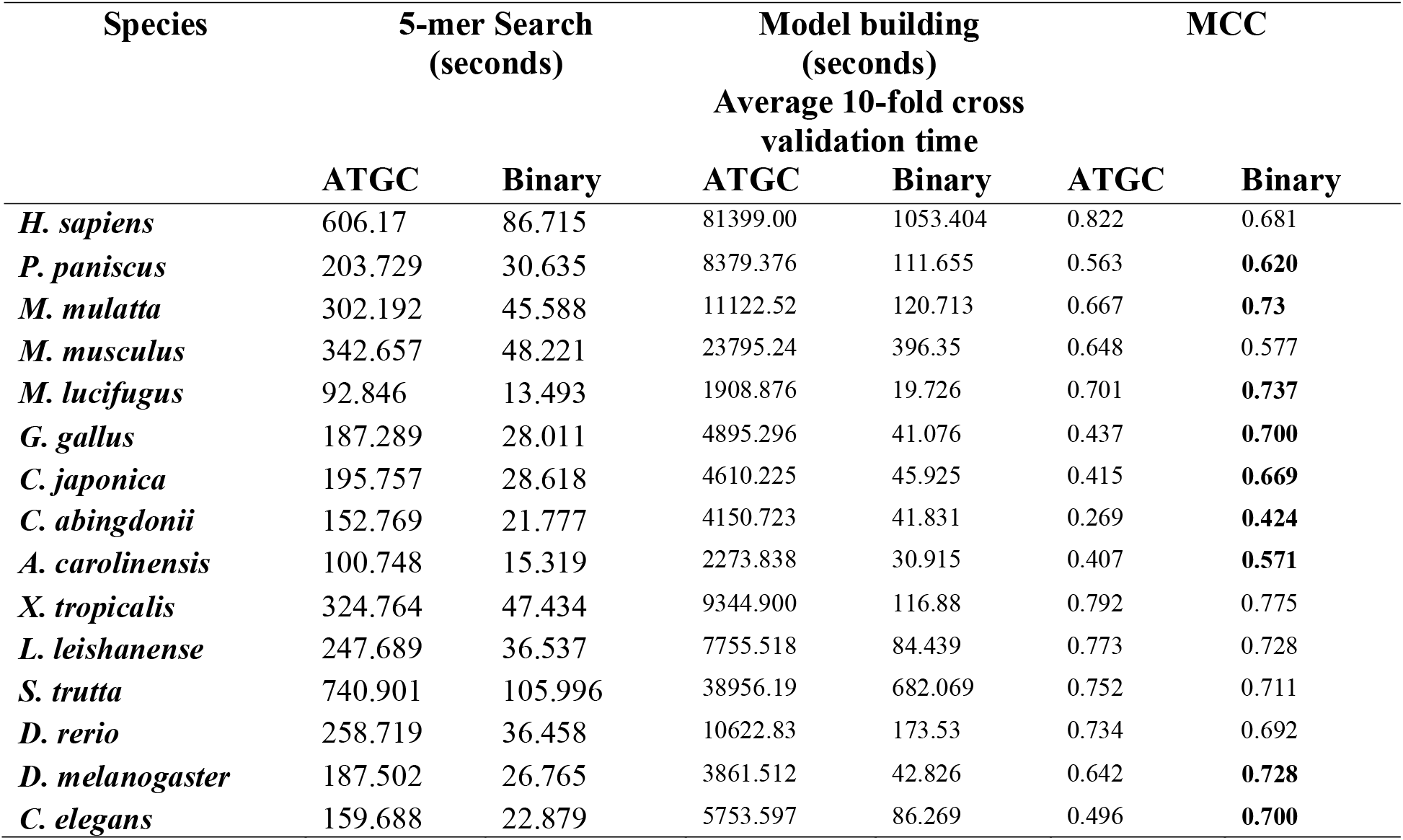
Comparison of time and MCC using traditional ATGC and binary 5-mer approach. Bold MCC presents species where the binary approach is superior to the ATGC approach.

### MCC scores correlates with evolutionary history among some species

Having established the success of the binary *k*-mer approach as an effective way to identify coding and non-coding sequences, we enhance this study by testing the ability of the ML model built by us. The ML model trained on human dataset was evaluated across several species such as *Pan paniscus*, *Gorilla gorilla*, *Mus musculus*, *Myotis lucifugus*, *Meleagris gallopavo*, *Podarcis muralis*, *Danio rerio*, *Ciona intestinalis* and *Saccharomyces cerevisiae*. As can be seen in Figure 5 (Supplementary Table S4), a ML model trained on humans gives high MCC for species closer to humans in the geological time scale. As humans diverge from other species in geological time scales, MCC score gradually decreases. For instance - MCC scores of reptiles (*P. muralis*: 0.27), fish (*D. rerio*: 0.27) and tunicates (*C. intestinalis*: 0.09) are lower than birds (*M. gallopavo*: 0.32) and mammals (*P. paniscus*: 0.55) which directly correlates the phylogenetic nearness of these species. It further emphasizes the fact that the model built on one species can be used to evaluate coding and non-coding transcripts of other species which are phylogenetically close to it. This data further attest to our binary approach where the MCC values correlated with the phylogenetic nearness of the species.

**Figure 5.**
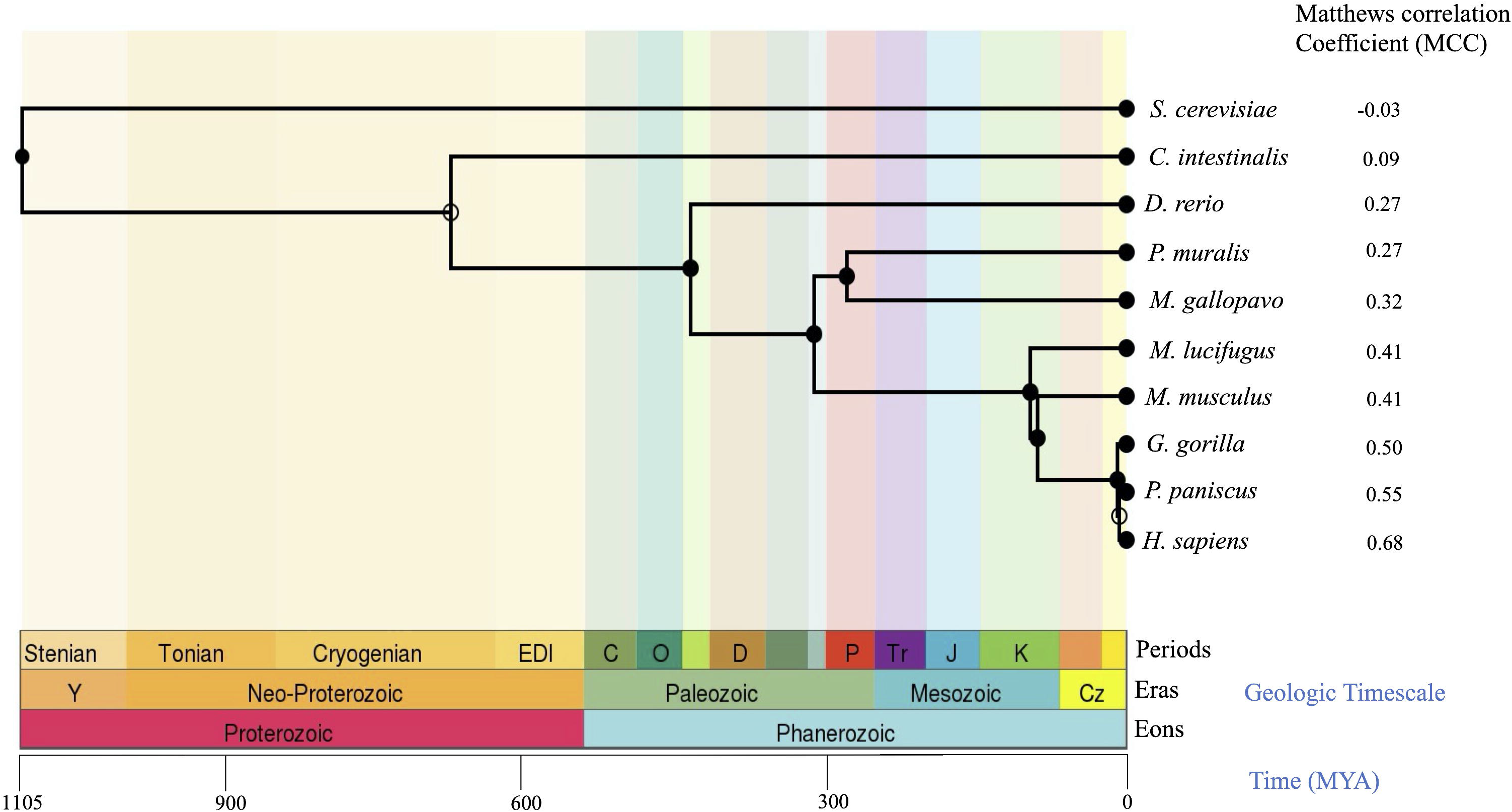
MCC scores correlates with evolutionary history among some species. The evolutionary geological time scale of various species. The MCC values of *k*NN (with 5 neighbours) model trained on *H.* sapiens and tested on other nine species correlates with their evolution. The species evolutionary closer to *H.* sapiens have higher MCC values as compared to those evolutionary farther to *H.* sapiens.

In summary, the major contributions of this paper are (i) distinguishing binary *k*-mer patterns were discovered, that were highly enriched in CDS and ncRNAs, (ii) a binary *k*-mer approach was devised to identify frequency of ‘binary signatures’ among CDS and ncRNAs thereby enabling us to use them as the features that effectively discriminate CDS and ncRNAs. This approach decreased the genome complexity as well as requirement of computer hardware to process the full genomic sequences consisting of four characters. Here, 2^k^ combinations were searched against the 4^k^ combinations of conventional ATGC *k*-mer approach, (iii) utilizing the complete data sets downloaded to build a robust ML model for classification. This approach utilizes all the crucial information in the CDS as well as ncRNAs as compared to the models built on a balanced dataset with only a few randomly chosen data samples and (iv) since next generation sequencing is generating huge amounts of data consisting of thousands of novel transcripts, our proposed method facilitates classification of novel CDS and ncRNAs. This can be achieved by building models of related known species and testing species that are phylogenetically closer to them.

## SUPPLEMENTARY DATA

Supplementary data are available online.

## AUTHOR CONTRIBUTIONS

VS conceived the idea. NP and PS performed the experiments. BK contributed to the ML setup and experiments. PA, SP, BK and VS supervised the study. VS wrote the first draft. PA, BK and VS edited and finalized the draft.

## ACKNOWLEDGEMENTS

NP and PS are recipients of PhD fellowship from UGC and CSIR respectively. VS is thankful to UGC for the Faculty Recharge award and start-up grant. We are thankful to Ishpreet Singh and Bhuvnesh Tanwar for their assistance in optimizing the python scripts.

