## Supplemental Table 1, Table 2, Table 3, Table 4 for "A novel binary *k*-mer approach for classification of coding and non-coding RNAs across diverse species"

**Table S1.** All possible patterns generated through binary *k*-mer approach for human CDS (*k* = 3) **Note:** This work was performed in all the fifteen species (*k* = 3 to *k* = 18). This is a representative data.

| **Pattern** | **Frequency** |
| --- | --- |
| 101 | 20447193 |
| 010 | 17861870 |
| 011 | 17040171 |
| 110 | 17038843 |
| 111 | 14784492 |
| 001 | 14477432 |
| 100 | 14406030 |
| 000 | 11751160 |

**Table S2.**  The *k*NN performance metric for all the fifteen species (*k* = 3 to *k* = 10)

1. *Homo Sapiens*

|  | ***k*-mer** | | | | | | | |
| --- | --- | --- | --- | --- | --- | --- | --- | --- |
|  | ***3*-mer** | ***4*-mer** | ***5*-mer** | ***6*-mer** | ***7*-mer** | ***8*-mer** | ***9*-mer** | ***10*-mer** |
| **Accuracy** | 0.795 | 0.826 | 0.858 | 0.875 | 0.884 | 0.89 | 0.891 | 0.89 |
| **F1 score** | 0.792 | 0.824 | 0.855 | 0.873 | 0.881 | 0.887 | 0.889 | 0.886 |
| **Precision** | 0.823 | 0.849 | 0.871 | 0.881 | 0.883 | 0.887 | 0.877 | 0.871 |
| **Recall** | 0.876 | 0.895 | 0.919 | 0.936 | 0.948 | 0.96 | 0.970 | 0.977 |
| **ROC-AUC** | 0.839 | 0.877 | 0.908 | 0.922 | 0.93 | 0.932 | 0.933 | 0.930 |
| **MCC** | 0.535 | 0.607 | 0.681 | 0.718 | 0.738 | 0.752 | 0.757 | 0.755 |

2. *Pan paniscus*

|  | *k*-mer | | | | | | | |
| --- | --- | --- | --- | --- | --- | --- | --- | --- |
|  | ***3*-mer** | ***4*-mer** | ***5*-mer** | ***6*-mer** | ***7*-mer** | ***8*-mer** | ***9*-mer** | ***10*-mer** |
| **Accuracy** | 0.881 | 0.895 | 0.897 | 0.89 | 0.881 | 0.872 | 0.864 | 0.857 |
| **F1 score** | 0.871 | 0.885 | 0.885 | 0.874 | 0.861 | 0.847 | 0.835 | 0.822 |
| **Precision** | 0.896 | 0.903 | 0.899 | 0.889 | 0.880 | 0.871 | 0.863 | 0.856 |
| **Recall** | 0.967 | 0.977 | 0.985 | 0.989 | 0.989 | 0.991 | 0.992 | 0.993 |
| **ROC-AUC** | 0.837 | 0.875 | 0.879 | 0.861 | 0.841 | 0.81 | 0.775 | 0.750 |
| **MCC** | 0.554 | 0.607 | 0.620 | 0.582 | 0.542 | 0.498 | 0.459 | 0.418 |

3. *Macaca mulatta*

|  | *k*-mer | | | | | | | |
| --- | --- | --- | --- | --- | --- | --- | --- | --- |
|  | ***3*-mer** | ***4*-mer** | ***5*-mer** | ***6*-mer** | ***7*-mer** | ***8*-mer** | ***9*-mer** | ***10*-mer** |
| **Accuracy** | 0.875 | 0.90 | 0.906 | 0.900 | 0.887 | 0.873 | 0.861 | 0.85 |
| **F1 score** | 0.869 | 0.895 | 0.902 | 0.895 | 0.880 | 0.864 | 0.849 | 0.834 |
| **Precision** | 0.890 | 0.908 | 0.910 | 0.904 | 0.893 | 0.881 | 0.868 | 0.857 |
| **Recall** | 0.955 | 0.968 | 0.974 | 0.974 | 0.97 | 0.966 | 0.965 | 0.966 |
| **ROC-AUC** | 0.875 | 0.91 | 0.919 | 0.910 | 0.894 | 0.873 | 0.851 | 0.826 |
| **MCC** | 0.624 | 0.702 | 0.73 | 0.704 | 0.662 | 0.615 | 0.574 | 0.531 |

4. *Mus musculus*

|  | *k*-mer | | | | | | | |
| --- | --- | --- | --- | --- | --- | --- | --- | --- |
|  | ***3*-mer** | ***4*-mer** | ***5*-mer** | ***6*-mer** | ***7*-mer** | ***8*-mer** | ***9*-mer** | ***10*-mer** |
| **Accuracy** | 0.810 | 0.832 | 0.854 | 0.863 | 0.867 | 0.865 | 0.864 | 0.859 |
| **F1 score** | 0.798 | 0.821 | 0.845 | 0.853 | 0.856 | 0.853 | 0.851 | 0.842 |
| **Precision** | 0.841 | 0.855 | 0.867 | 0.870 | 0.870 | 0.865 | 0.861 | 0.852 |
| **Recall** | 0.924 | 0.935 | 0.953 | 0.962 | 0.968 | 0.973 | 0.979 | 0.984 |
| **ROC-AUC** | 0.787 | 0.827 | 0.857 | 0.867 | 0.872 | 0.867 | 0.860 | 0.850 |
| **MCC** | 0.439 | 0.508 | 0.577 | 0.602 | 0.611 | 0.609 | 0.605 | 0.589 |

5. *Myotis lucifugus*

|  | *k*-mer | | | | | | | |
| --- | --- | --- | --- | --- | --- | --- | --- | --- |
|  | ***3*-mer** | ***4*-mer** | ***5*-mer** | ***6*-mer** | ***7*-mer** | ***8*-mer** | ***9*-mer** | ***10*-mer** |
| **Accuracy** | 0.911 | 0.935 | 0.931 | 0.918 | 0.908 | 0.90 | 0.894 | 0.890 |
| **F1 score** | 0.904 | 0.930 | 0.925 | 0.907 | 0.894 | 0.883 | 0.874 | 0.869 |
| **Precision** | 0.917 | 0.933 | 0.925 | 0.91 | 0.90 | 0.892 | 0.886 | 0.883 |
| **Recall** | 0.981 | 0.993 | 0.998 | 0.999 | 0.999 | 1 | 1 | 1 |
| **ROC-AUC** | 0.874 | 0.929 | 0.922 | 0.888 | 0.846 | 0.809 | 0.783 | 0.766 |
| **MCC** | 0.666 | 0.763 | 0.737 | 0.694 | 0.649 | 0.617 | 0.588 | 0.575 |

6. *Gallus gallus*

|  | *k*-mer | | | | | | | |
| --- | --- | --- | --- | --- | --- | --- | --- | --- |
|  | ***3*-mer** | ***4*-mer** | ***5*-mer** | ***6*-mer** | ***7*-mer** | ***8*-mer** | ***9*-mer** | ***10*-mer** |
| **Accuracy** | 0.863 | 0.879 | 0.885 | 0.885 | 0.876 | 0.86 | 0.835 | 0.808 |
| **F1 score** | 0.858 | 0.875 | 0.880 | 0.878 | 0.866 | 0.845 | 0.809 | 0.769 |
| **Precision** | 0.879 | 0.890 | 0.89 | 0.882 | 0.868 | 0.847 | 0.820 | 0.794 |
| **Recall** | 0.942 | 0.952 | 0.962 | 0.973 | 0.98 | 0.987 | 0.991 | 0.994 |
| **ROC-AUC** | 0.874 | 0.902 | 0.906 | 0.908 | 0.900 | 0.882 | 0.85 | 0.808 |
| **MCC** | 0.633 | 0.68 | 0.700 | 0.693 | 0.668 | 0.624 | 0.550 | 0.471 |

7. *Coturnix japonica*

|  | *k*-mer | | | | | | | |
| --- | --- | --- | --- | --- | --- | --- | --- | --- |
|  | ***3*-mer** | ***4*-mer** | ***5*-mer** | ***6*-mer** | ***7*-mer** | ***8*-mer** | ***9*-mer** | ***10*-mer** |
| **Accuracy** | 0.863 | 0.880 | 0.881 | 0.872 | 0.862 | 0.845 | 0.828 | 0.815 |
| **F1 score** | 0.858 | 0.875 | 0.874 | 0.861 | 0.847 | 0.822 | 0.798 | 0.776 |
| **Precision** | 0.883 | 0.89 | 0.882 | 0.867 | 0.853 | 0.834 | 0.817 | 0.804 |
| **Recall** | 0.941 | 0.958 | 0.971 | 0.978 | 0.984 | 0.988 | 0.991 | 0.993 |
| **ROC-AUC** | 0.869 | 0.898 | 0.897 | 0.888 | 0.871 | 0.849 | 0.816 | 0.790 |
| **MCC** | 0.623 | 0.669 | 0.668 | 0.644 | 0.615 | 0.561 | 0.509 | 0.459 |

8. *Chelonoidis abingdonii*

|  | *k*-mer | | | | | | | |
| --- | --- | --- | --- | --- | --- | --- | --- | --- |
|  | ***3*-mer** | ***4*-mer** | ***5*-mer** | ***6*-mer** | ***7*-mer** | ***8*-mer** | ***9*-mer** | ***10*-mer** |
| **Accuracy** | 0.929 | 0.936 | 0.932 | 0.924 | 0.921 | 0.920 | 0.918 | 0.92 |
| **F1 score** | 0.915 | 0.921 | 0.912 | 0.896 | 0.888 | 0.887 | 0.883 | 0.885 |
| **Precision** | 0.937 | 0.939 | 0.932 | 0.924 | 0.921 | 0.920 | 0.918 | 0.92 |
| **Recall** | 0.989 | 0.995 | 0.998 | 1 | 1 | 1 | 1 | 1 |
| **ROC-AUC** | 0.770 | 0.813 | 0.783 | 0.709 | 0.634 | 0.598 | 0.58 | 0.574 |
| **MCC** | 0.415 | 0.475 | 0.424 | 0.314 | 0.242 | 0.221 | 0.200 | 0.199 |

9. *Anolis carolinensis*

|  | *k*-mer | | | | | | | |
| --- | --- | --- | --- | --- | --- | --- | --- | --- |
|  | ***3*-mer** | ***4*-mer** | ***5*-mer** | ***6*-mer** | ***7*-mer** | ***8*-mer** | ***9*-mer** | ***10*-mer** |
| **Accuracy** | 0.809 | 0.836 | 0.835 | 0.816 | 0.799 | 0.786 | 0.778 | 0.770 |
| **F1 score** | 0.801 | 0.826 | 0.820 | 0.794 | 0.768 | 0.749 | 0.734 | 0.721 |
| **Precision** | 0.832 | 0.842 | 0.83 | 0.807 | 0.789 | 0.777 | 0.768 | 0.761 |
| **Recall** | 0.916 | 0.946 | 0.966 | 0.974 | 0.978 | 0.98 | 0.983 | 0.986 |
| **ROC-AUC** | 0.814 | 0.856 | 0.86 | 0.847 | 0.816 | 0.779 | 0.750 | 0.722 |
| **MCC** | 0.51 | 0.578 | 0.571 | 0.523 | 0.474 | 0.433 | 0.407 | 0.384 |

10. *Xenopus tropicalis*

|  | *k*-mer | | | | | | | |
| --- | --- | --- | --- | --- | --- | --- | --- | --- |
|  | ***3*-mer** | ***4*-mer** | ***5*-mer** | ***6*-mer** | ***7*-mer** | ***8*-mer** | ***9*-mer** | ***10*-mer** |
| **Accuracy** | 0.987 | 0.99 | 0.990 | 0.990 | 0.99 | 0.99 | 0.99 | 0.99 |
| **F1 score** | 0.986 | 0.989 | 0.989 | 0.989 | 0.988 | 0.99 | 0.99 | 0.988 |
| **Precision** | 0.989 | 0.990 | 0.991 | 0.990 | 0.99 | 0.99 | 0.99 | 0.99 |
| **Recall** | 0.998 | 1 | 1 | 1 | 1 | 1 | 1 | 1 |
| **ROC-AUC** | 0.881 | 0.902 | 0.896 | 0.88 | 0.857 | 0.846 | 0.838 | 0.835 |
| **MCC** | 0.699 | 0.754 | 0.775 | 0.766 | 0.762 | 0.762 | 0.763 | 0.761 |

11. *Leptobrachium leishanense*

|  | *k*-mer | | | | | | | |
| --- | --- | --- | --- | --- | --- | --- | --- | --- |
|  | ***3*-mer** | ***4*-mer** | ***5*-mer** | ***6*-mer** | ***7*-mer** | ***8*-mer** | ***9*-mer** | ***10*-mer** |
| **Accuracy** | 0.987 | 0.99 | 0.992 | 0.992 | 0.992 | 0.992 | 0.992 | 0.992 |
| **F1 score** | 0.985 | 0.989 | 0.990 | 0.991 | 0.991 | 0.991 | 0.991 | 0.991 |
| **Precision** | 0.989 | 0.991 | 0.992 | 0.993 | 0.993 | 0.992 | 0.992 | 0.992 |
| **Recall** | 0.998 | 0.998 | 1 | 1 | 1 | 1 | 1 | 1 |
| **ROC-AUC** | 0.821 | 0.865 | 0.881 | 0.868 | 0.858 | 0.843 | 0.842 | 0.839 |
| **MCC** | 0.561 | 0.668 | 0.728 | 0.742 | 0.752 | 0.743 | 0.747 | 0.745 |

12. *Salmo trutta*

|  | *k*-mer | | | | | | | |
| --- | --- | --- | --- | --- | --- | --- | --- | --- |
|  | ***3*-mer** | ***4*-mer** | ***5*-mer** | ***6*-mer** | ***7*-mer** | ***8*-mer** | ***9*-mer** | ***10*-mer** |
| **Accuracy** | 0.973 | 0.976 | 0.978 | 0.979 | 0.979 | 0.979 | 0.979 | 0.979 |
| **F1 score** | 0.971 | 0.974 | 0.977 | 0.977 | 0.978 | 0.977 | 0.977 | 0.977 |
| **Precision** | 0.979 | 0.982 | 0.983 | 0.983 | 0.984 | 0.984 | 0.983 | 0.983 |
| **Recall** | 0.993 | 0.994 | 0.995 | 0.995 | 0.995 | 0.995 | 0.995 | 0.995 |
| **ROC-AUC** | 0.879 | 0.901 | 0.91 | 0.916 | 0.918 | 0.919 | 0.915 | 0.912 |
| **MCC** | 0.631 | 0.675 | 0.711 | 0.718 | 0.717 | 0.717 | 0.717 | 0.713 |

13. *Danio rerio*

|  | *k*-mer | | | | | | | |
| --- | --- | --- | --- | --- | --- | --- | --- | --- |
|  | ***3*-mer** | ***4*-mer** | ***5*-mer** | ***6*-mer** | ***7*-mer** | ***8*-mer** | ***9*-mer** | ***10*-mer** |
| **Accuracy** | 0.905 | 0.925 | 0.934 | 0.937 | 0.938 | 0.935 | 0.934 | 0.931 |
| **F1 score** | 0.897 | 0.92 | 0.929 | 0.932 | 0.934 | 0.93 | 0.928 | 0.924 |
| **Precision** | 0.924 | 0.937 | 0.942 | 0.944 | 0.944 | 0.941 | 0.938 | 0.935 |
| **Recall** | 0.970 | 0.979 | 0.984 | 0.986 | 0.987 | 0.988 | 0.988 | 0.989 |
| **ROC-AUC** | 0.838 | 0.881 | 0.896 | 0.898 | 0.893 | 0.881 | 0.872 | 0.861 |
| **MCC** | 0.543 | 0.647 | 0.692 | 0.704 | 0.711 | 0.697 | 0.687 | 0.671 |

14. *Drosophila melanogaster*

|  | *k*-mer | | | | | | | |
| --- | --- | --- | --- | --- | --- | --- | --- | --- |
|  | ***3*-mer** | ***4*-mer** | ***5*-mer** | ***6*-mer** | ***7*-mer** | ***8*-mer** | ***9*-mer** | ***10*-mer** |
| **Accuracy** | 0.943 | 0.949 | 0.949 | 0.95 | 0.948 | 0.944 | 0.942 | 0.937 |
| **F1 score** | 0.940 | 0.946 | 0.946 | 0.945 | 0.942 | 0.938 | 0.934 | 0.928 |
| **Precision** | 0.956 | 0.958 | 0.955 | 0.953 | 0.949 | 0.945 | 0.942 | 0.934 |
| **Recall** | 0.980 | 0.986 | 0.989 | 0.992 | 0.994 | 0.995 | 0.996 | 0.996 |
| **ROC-AUC** | 0.908 | 0.917 | 0.915 | 0.907 | 0.891 | 0.877 | 0.861 | 0.835 |
| **MCC** | 0.699 | 0.726 | 0.728 | 0.728 | 0.711 | 0.692 | 0.676 | 0.646 |

15. *Caenorhabditis elegans*

|  | *k*-mer | | | | | | | |
| --- | --- | --- | --- | --- | --- | --- | --- | --- |
|  | ***3*-mer** | ***4*-mer** | ***5*-mer** | ***6*-mer** | ***7*-mer** | ***8*-mer** | ***9*-mer** | ***10*-mer** |
| **Accuracy** | 0.905 | 0.914 | 0.903 | 0.879 | 0.857 | 0.841 | 0.834 | 0.829 |
| **F1 score** | 0.900 | 0.909 | 0.893 | 0.862 | 0.830 | 0.808 | 0.797 | 0.790 |
| **Precision** | 0.914 | 0.912 | 0.896 | 0.87 | 0.848 | 0.835 | 0.831 | 0.827 |
| **Recall** | 0.97 | 0.981 | 0.991 | 0.995 | 0.995 | 0.993 | 0.99 | 0.988 |
| **ROC-AUC** | 0.898 | 0.919 | 0.905 | 0.852 | 0.791 | 0.732 | 0.699 | 0.683 |
| **MCC** | 0.706 | 0.732 | 0.700 | 0.618 | 0.535 | 0.469 | 0.429 | 0.407 |

**Table S3.** Comparison of conventional ATGC and binary *5*-mer approach. Bold MCC values represents the case where binary approach performs better than conventional ATGC approach

| **Species** | ***5*-mer Search**  **(seconds)** | | ***5*-mer Model (seconds)** | | **Accuracy** | | **F1 Score** | | **Precision** | | **Recall** | | **ROC-AUC** | | **MCC** | |
| --- | --- | --- | --- | --- | --- | --- | --- | --- | --- | --- | --- | --- | --- | --- | --- | --- |
|  | **ATGC** | **Binary** | **ATGC** | **Binary** | **ATGC** | **Binary** | **ATGC** | **Binary** | **ATGC** | **Binary** | **ATGC** | **Binary** | **ATGC** | **Binary** | **ATGC** | **Binary** |
| *H. sapiens* | 606.17 | 86.715 | 81399.00 | 1053.404 | 0.919 | 0.858 | 0.91 | 0.855 | 0.91 | 0.871 | 1 | 0.919 | 0.919 | 0.908 | 0.822 | 0.681 |
| *P. paniscus* | 203.729 | 30.635 | 8379.376 | 111.655 | 0.885 | 0.897 | 0.865 | 0.885 | 0.882 | 0.899 | 0.992 | 0.985 | 0.830 | 0.879 | 0.563 | **0.620** |
| *M. mulatta* | 302.192 | 45.588 | 11122.52 | 120.713 | 0.888 | 0.906 | 0.882 | 0.902 | 0.895 | 0.910 | 0.967 | 0.974 | 0.895 | 0.919 | 0.667 | **0.73** |
| *M. musculus* | 342.657 | 48.221 | 23795.24 | 396.35 | 0.877 | 0.854 | 0.866 | 0.845 | 0.872 | 0.867 | 0.982 | 0.953 | 0.886 | 0.857 | 0.648 | 0.577 |
| *M. lucifugus* | 92.846 | 13.493 | 1908.876 | 19.726 | 0.919 | 0.931 | 0.909 | 0.925 | 0.910 | 0.925 | 1 | 0.998 | 0.837 | 0.922 | 0.701 | **0.737** |
| *G. gallus* | 187.289 | 28.011 | 4895.296 | 41.076 | 0.799 | 0.885 | 0.754 | 0.880 | 0.787 | 0.89 | 0.996 | 0.962 | 0.805 | 0.906 | 0.437 | **0.700** |
| *C. japonica* | 195.757 | 28.618 | 4610.225 | 45.925 | 0.806 | 0.881 | 0.760 | 0.874 | 0.797 | 0.882 | 0.993 | 0.971 | 0.785 | 0.897 | 0.415 | **0.668** |
| *C. abingdonii* | 152.769 | 21.777 | 4150.723 | 41.831 | 0.921 | 0.932 | 0.89 | 0.912 | 0.921 | 0.932 | 1 | 0.998 | 0.592 | 0.783 | 0.269 | **0.424** |
| *A. carolinensis* | 100.748 | 15.319 | 2273.838 | 30.915 | 0.778 | 0.835 | 0.729 | 0.820 | 0.766 | 0.83 | 0.99 | 0.966 | 0.756 | 0.86 | 0.407 | **0.571** |
| *X. tropicalis* | 324.764 | 47.434 | 9344.900 | 116.88 | 0.991 | 0.990 | 0.990 | 0.989 | 0.990 | 0.991 | 1 | 1 | 0.863 | 0.896 | 0.792 | 0.775 |
| *L. leishanense* | 247.689 | 36.537 | 7755.518 | 84.439 | 0.992 | 0.992 | 0.992 | 0.990 | 0.993 | 0.992 | 1 | 1 | 0.854 | 0.881 | 0.773 | 0.728 |
| *S. trutta* | 740.901 | 105.996 | 38956.19 | 682.069 | 0.982 | 0.978 | 0.980 | 0.977 | 0.985 | 0.983 | 0.996 | 0.995 | 0.927 | 0.91 | 0.752 | 0.711 |
| *D. rerio* | 258.719 | 36.458 | 10622.83 | 173.53 | 0.943 | 0.934 | 0.939 | 0.929 | 0.946 | 0.942 | 0.992 | 0.984 | 0.891 | 0.896 | 0.734 | 0.692 |
| *D. melanogaster* | 187.502 | 26.765 | 3861.512 | 42.826 | 0.937 | 0.949 | 0.928 | 0.946 | 0.937 | 0.955 | 0.997 | 0.989 | 0.856 | 0.915 | 0.642 | **0.728** |
| *C. elegans* | 159.688 | 22.879 | 5753.597 | 86.269 | 0.848 | 0.903 | 0.817 | 0.893 | 0.840 | 0.896 | 0.994 | 0.991 | 0.725 | 0.905 | 0.496 | **0.700** |

**Table S4**. Validation of proposed novel binary *k*NN model trained on *Homo sapiens* and tested on nine diverse species.

|  | **Species tested on model trained on *Homo sapiens*** | | | | | | | | | |
| --- | --- | --- | --- | --- | --- | --- | --- | --- | --- | --- |
|  | ***H. sapiens*** | ***P. paniscus*** | ***G. gorilla*** | ***M. musculus*** | ***M. lucifugus*** | ***M. gallopavo*** | ***P. muralis*** | ***D. rerio*** | ***C. intestinalis*** | ***S. cerevisiae*** |
| **Accuracy** | 0.858 | 0.871 | 0.876 | 0.784 | 0.827 | 0.862 | 0.875 | 0.788 | 0.798 | 0.686 |
| **F1 score** | 0.855 | 0.922 | 0.927 | 0.858 | 0.858 | 0.923 | 0.932 | 0.871 | 0.886 | 0.811 |
| **Precision** | 0.871 | 0.914 | 0.924 | 0.853 | 0.853 | 0.961 | 0.973 | 0.916 | 0.982 | 0.936 |
| **Recall** | 0.919 | 0.93 | 0.930 | 0.863 | 0.863 | 0.889 | 0.894 | 0.830 | 0.807 | 0.715 |
| **ROC-AUC** | 0.908 | 0.767 | 0.749 | 0.702 | 0.702 | 0.715 | 0.71 | 0.671 | 0.612 | 0.475 |
| **MCC** | 0.681 | 0.552 | 0.505 | 0.411 | 0.410 | 0.321 | 0.274 | 0.273 | 0.09 | -0.03 |
